# BLOBFISH: Bipartite Limited Subnetworks from Multiple Observations using Breadth-First Search with Constrained Hops

**DOI:** 10.1101/2025.03.11.642378

**Authors:** Tara Eicher, Marouen Ben Guebila, John Quackenbush

**Affiliations:** Department of Biostatistics, T.H. Chan School of Public Health, Harvard University, Boston, MA, 02115, USA; Channing Division of Network Medicine, Brigham and Women’s Hospital, Boston, MA, 02115, USA

**Author notes:** **Availability and Implementation** Source code is available from GitHub as part of the *netZooR* R package (v1.6) (https://github.com/netZoo/netZooR) (Ben Guebila, et al., 2023).

## Abstract

**Summary:** In analyzing gene regulatory network models, a common question is how members of a particular set of genes are connected. For example, one might want to explore network relationship between a set of differentially expressed genes, a gene set previously reported in the literature, or elements of one or more pathways. BLOBFISH uses a breadth-first search algorithm adapted to bipartite graphs to identify a compact subnetwork connecting the members of a pre-specified set of genes, providing a regulatory context that can shed light on specific mechanisms involved in a phenotype and its development. We demonstrate the use of BLOBFISH to extract gene regulatory subnetworks reflecting tissue specificity using publicly available data from the Genotype Tissue Expression (GTEx) project.

## Introduction

It is common to think of biological states as being defined by specific genetic variants or patterns of expression, but we have come to recognize that the real drivers of phenotypes are complex networks of interacting cellular elements that drive the processes specific to each phenotype and that characterize the transitions between states. There are many tools for inferring networks that represent specific aspects of biological systems and these have proven their value in characterizing the relationships between genes and other entities in the context of understanding phenotypic differences (Eicher, et al., 2020; Weighill, et al., 2021). Although unipartite networks such as gene co-expression based on correlations or partial correlations are frequently used (Langfelder and Horvath, 2008), bipartite networks provide the opportunity to identify interactions between different types of cellular elements, including genes and their regulators. Examples of bipartite networks include expression quantitative trait loci (eQTL) networks (Fagny, et al., 2017; Platig, et al., 2016), gene regulatory networks (GRNs) (Glass, et al., 2013; Glass, et al., 2013; Kuijjer, et al., 2020), and multi-omic partial correlation networks (Eicher, et al., 2023; Shutta, et al., 2023).

Although networks can capture genome-scale interactions, interpreting such results can be challenging, particularly in the context of exploring a hypothesis regarding a subset of genes thought to be relevant to a phenotype under study (Edge, et al., 2019). Interest in examining network relationships between a subset of “interesting” genes has motivated the development of tools to evaluate connectivity between a set of “seed” nodes, including methods based on random walks with restarts (Chakrabarty, et al., 2021; Liao, et al., 2020) and Steiner trees (White and Ma’ayan, 2007). However, most subgraph extracting methods have a stochastic component so that multiple applications to the same starting network, or changes in how the network is recorded, may lead to different results. There are also deterministic methods that can identify unique paths connecting seed nodes, including some based on depth-first traversal (Berger, et al., 2007) and shortest paths (Ho, et al., 2012; Keane, et al., 2015), but in applications such as drug targeting and pathway analysis, where biological systems often have redundancies, deterministic methods can miss multiple connecting paths that define more biologically relevant patterns of interactivity (Lee, et al., 2023; Ogris, et al., 2022).

A key hypothesis in network analysis is that networks not only differ between phenotypes but also vary among individuals within any specific phenotypic group. Methods such LIONESS (Kuijjer, et al., 2019), BONOBO (Saha, et al., 2023), and SWEET (Chen, et al., 2023), used in conjunction with other methods, can infer networks for each individual in a population and there are a number of methods that can be used to for single-cell specific network inference (such as those reviewed by Pratapa et al. (Pratapa, et al., 2020)). This suggests that a useful strategy in mapping subnetworks connecting genes would be to identify robust edges that are consistent across individuals, providing greater confidence in the identified connectivity between genes.

## Approach

We developed BLOBFISH (**B**ipartite **L**imited Subnetworks from Multiple **O**bservations using **B**readth-**Fi**rst **S**earch with Constrained **H**ops) to find a robust subnetwork connecting a seed set of nodes in collections of individual sample bipartite networks to find patterns of connection consistent across observations. Conceptually, BLOBFISH is based on the idea that each node in a bipartite graph has a sphere of influence that can be obtained using breadth-first search with a user-defined constraint on the number of allowed hops between nodes such that the connectivity between two nodes represents paths to shared nodes within each node’s sphere of influence. This allows multiple paths to connect individual nodes in a way that mirrors the redundancy and shared functionality of genes known to exist in biological networks. To provide confidence in the resulting subnetworks, BLOBFISH requires that all edges in the subnetwork must have statistically significant weights when compared against a null model. Because BLOBFISH operates on bipartite graphs, all connecting paths can be found by evaluating only shared nodes equidistant from a pair of seed nodes (see Theorem 1 in Supplementary File 1). By making use of this property in addition to being deterministic, BLOBFISH provides runtime advantages over a naïve exploration of all possible paths across spheres of influence. The BLOBFISH algorithm is described as pseudocode in Supplementary File 1 together with analysis of some of its performance characteristics.

BLOBFISH takes as input a collection of gene regulatory networks derived by applying a specific GRN inference method to an experimental dataset and a null network distribution by using multiple randomizations of the expression levels (or other biological measurements) and using the same network inference method to generate a collection of network models. BLOBFISH analyzes the collection of individual-sample network graphs and, for a set of seed genes, and first uses the Wilcoxon rank-sum test to estimate the significance of the inferred regulatory edges by comparing these to the null network; edges that are significant (based on a user-defined sphere of influence parameter, α, set by default to 2) are retained in a reduced network model. BLOBFISH uses a breadth-first search (BFS) to identify neighborhoods for each gene in the reduced network. BFS explores gene nodes and regulatory edges connecting them systematically starting from each of the seed genes. It explores all neighbors of the current node before moving to the next level of neighbors, ensuring that all nodes are visited in order of their distance from the source, resulting in a sphere of influence for each gene based on a user-determined parameter. The algorithm then finds all possible paths connecting the genes based on the overlap of the collective spheres of influence to create more or more subnetworks connecting the seed gene set.

BLOBFISH is integrated into the *netZooR* R package (v1.6) (https://github.com/netZoo/netZooR) (Ben Guebila, et al., 2023) and can be run using a call to the function *RunBLOBFISH*(). The additional functions *PlotNetwork*() and *GenerateNullPANDADistribution*() support plotting the subnetwork and generating the null distribution for bipartite networks generated using PANDA’s message-passing framework or those of its related methods (Glass, et al., 2013; Kuijjer, et al., 2020; Osorio, et al., 2024; Sonawane, et al., 2021; Weighill, et al., 2022); the latter function could be easily modified to use other network inference methods. BLOBFISH depends on the R packages *igraph* and *matrixTests*. The script used in our analysis is available from https://github.com/QuackenbushLab/BLOBFISH_paper_scripts.

## Results

As a test of BLOBFISH, we downloaded from the GRAND database (Ben Guebila, et al., 2022) GRN models that had been inferred by applying PANDA+LIONESS to GTEx data (Lonsdale, et al., 2013; Lopes-Ramos, et al., 2020) for five tissues (subcutaneous adipose, skeletal muscle, lung tissue, skin, and aorta). To minimize influence of age and biological sex on the gene regulatory networks, we used only networks generated for males between the ages of 20-29, resulting in *n* = 18 for subcutaneous adipose, *n* = 27 for skeletal muscle, *n* = 15 for lung tissue, *n* = 31 for skin, and *n* = 15 for aorta. We used 26 genes reported to be typical of skeletal muscle by Bortoluzzi et al (Bortoluzzi, et al., 2000) and seven genes involved in adipogenesis (Ou-yang and Dai, 2023) as seed gene sets (available in Supplementary Tables 1 and 2).

We ran BLOBFISH using both the skeletal muscle and adipogenesis seed gene sets on each tissue-specific collection of bipartite gene regulatory networks using a null distribution generated using *GenerateNullPANDADistribution*(), α = 0.05, and a hop constraint of 2 (restricting paths between seed genes to contain a single transcription factor; full subnetworks can be found in Supplementary Tables 3-7). This provides both positive and negative controls that can be used to assess BLOBFISH’s performance. To evaluate only the tissue-specific connectivity between genes in the seed set and exclude connectivity shared between tissues, we retained only edges that were exclusive to each tissue type’s subnetwork. The results can be found in Supplementary Tables 8-12.

To assess the significance of the connectivity we found among gene sets associated with skeletal muscle development and adipogenesis in the relevant tissues, we used BLOBFISH to assess these and 100 random sets of 33 (26 + 7) genes in all five tissues for which we had network models.

We expect that few if any genes are expressed in only a single tissue and many genes are expressed in a large number of tissues; indeed, for each set of seed genes, we found subnetworks linking some subset of those seed genes in each of the five tissues in nearly every iteration. But we also expect phenotype-specific sets of genes to be highly connected in the relevant tissue since the some level of coordinated regulation of these genes defines, or is defined by, that tissue. Consistent with this, the mean count of transcription factors co-regulating each pair of skeletal-muscle-associated genes was highest in the skeletal-muscle-specific subnetwork and that the mean count of transcription factors co-regulating each pair of adipogenesis-associated genes was highest in the subcutaneous-adipose-specific subnetwork (Figure 1 and Table 1; see Supplementary Figure 1 for the other tissue-specific subnetworks). Further, in the relevant tissue, the mean counts for tissue-specific gene sets were considerably higher (with statistically significant Z-scores) than the mean counts obtained from the random gene sets. In contrast, the transcription factor connectivity for the same adipose and muscle gene sets were essentially at background levels in lung, skin, and aorta based on the distributions for random gene sets. This suggests that BLOBFISH can extract meaningful regulatory subnetwork connections between a tissue- or disease-specific set of genes when applied to gene regulatory networks inferred in a relevant tissue using PANDA+LIONESS.

**Figure 1.**
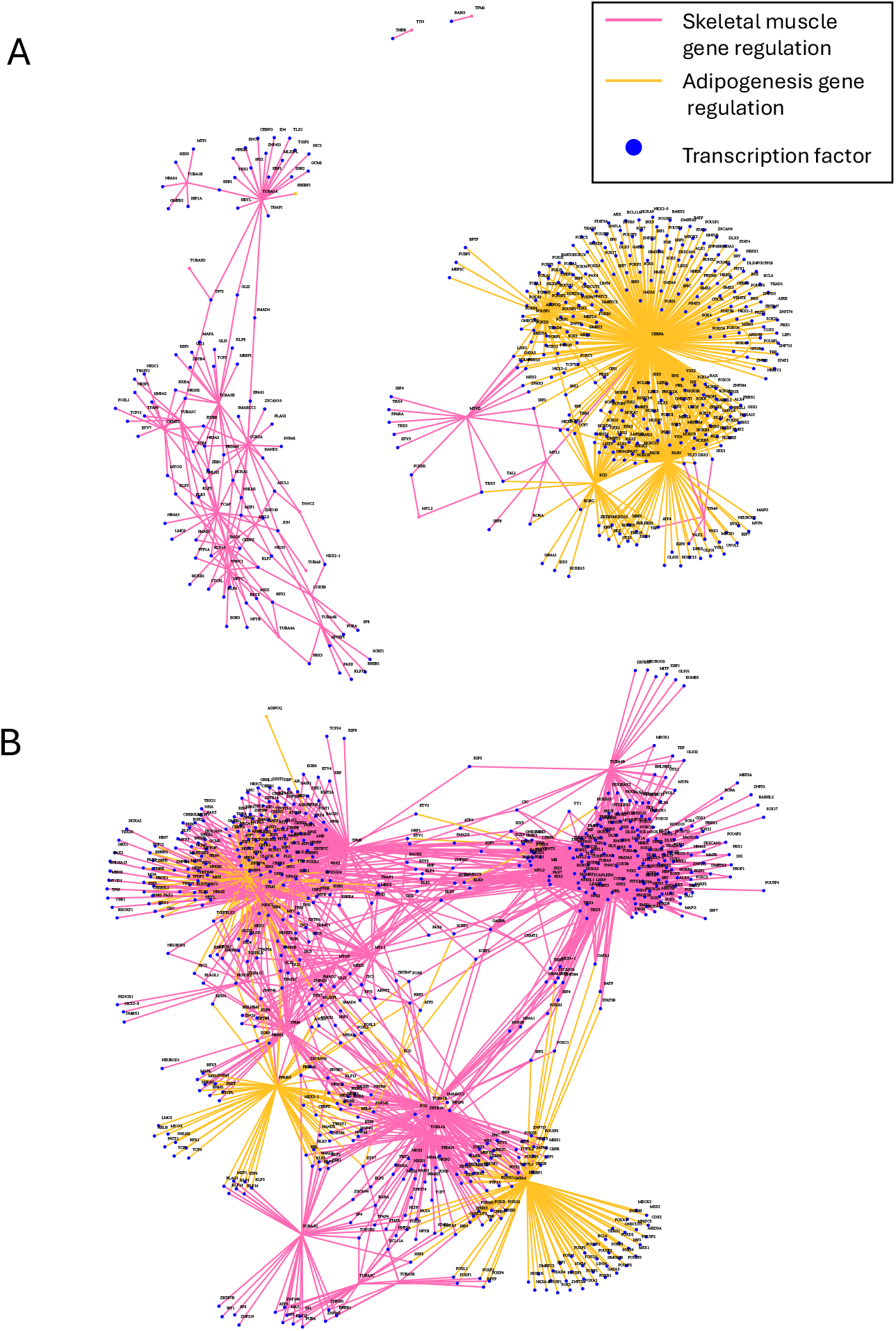
Tissue-specific gene regulatory subnetworks for GTEx skeletal muscle samples for males 20-29 in (A) subcutaneous adipose and (B) skeletal muscle reveal more co-regulatory connections between skeletal-muscle-associated genes (pink) in skeletal muscle and more co-regulatory connections between adipogenesis-associated genes (gold) in subcutaneous adipose.

**Table 1.** The mean number of transcription factors co-regulating skeletal-muscle-associated and adipogenesis-associated genes in each tissue, compared to the mean number of transcription factors co-regulating random subsets of genes.

|  | Subcutaneous<br>adipose | Skeletal<br>muscle | Lung tissue | Skin | Aorta |
| --- | --- | --- | --- | --- | --- |
| $\mu_{\text{rand}}^1 (\sigma^2)$ | 0.48 (0.35) | 6.96 (2.18) | 0.42 (0.24) | 1.14 (0.59) | 0.25 (0.20) |
| $\mu_{\text{skel}} (\text{z-score}^3)$ | 0.35 (-0.37) | **12.50 (2.54)** | 0.51 (0.38) | 0.26 (-1.49) | 0.10 (-0.75) |
| $\mu_{\text{adip}} (\text{z-score})$ | **9.10 (24.63)** | 0.48 (-2.97) | 0.00 (-1.75) | 1.00 (-0.24) | 0.43 (0.90) |

<sup>1</sup> Mean shared transcription factor counts for random (*rand*), skeletal-muscle-associated (*skel*), and adipogenesis-associated (*adip*) gene sets, respectively.
<sup>2</sup> Standard deviation of shared transcription factor counts across random gene sets.
<sup>3</sup> z-score when computed against $\mu_{\text{rand}}$ and $\sigma$ .

Beyond the tissue specificity we found for connected subnetworks of tissue-relevant gene sets, one would expect that the additional genes connected to the seed set found by BLOBFISH might further help explain the phenotype that the input gene set was meant to capture. To test this, we took the BLOBFISH subnetworks for each tissue generated using the combined skeletal muscle and adipogenesis genes as seeds, and for each performed pre-ranked gene set enrichment analysis on the transcription factors and genes using the *fgsea* R package (Korotkevich, et al., 2021) with *FDR*-adjusted *p*-value < 0.05 and gene sets from the Molecular signatures Database (MSigDB, version 2023.2) (Liberzon, et al., 2015; Subramanian, et al., 2005); the number of edges adjacent to each transcription factor or gene was used as the input ranking score. We then compared these results to pathways enriched in each tissue using the log_2_(fold change) of the expression levels of the seed as input.

The only functional class that was enriched in the subcutaneous adipose tissue subnetwork was adipogenesis, but this was not surprising given the small number of adipogenesis-relevant genes in the input (seven). In comparison, pathway analysis using expression of the seed resulted in enrichment of PPAR signaling, one step in adipogenesis (Ghaben and Scherer, 2019). In the skeletal muscle network subnetwork seeded by twenty-six genes, we found enrichment for both muscle contraction and Parkinson’s Disease pathways, whereas no pathways were enriched using gene expression data for this the seed set. Impaired skeletal muscle function is a hallmark of Parkinson’s Disease (Murphy and Lynch, 2023), this result provides further evidence that BLOBFISH can use tissue-specific gene regulatory networks to aggregate sets of genes linked through common regulators that share a common biological function. The results for all tissues are presented in Supplementary Tables 13-17 (for network-based analysis) and 18-21 (for expression-based analysis).

## Discussion

A frequent question in the analysis of any biologically interesting phenotype is how subsets of genes previously identified as relevant interact with each other within the context of the regulatory processes that ultimately define each biological state. Methods that attempt to connect genes within those subsets, including methods using correlation analysis, fail to provide information on the broader regulatory context in which these genes exist. Genome-wide gene regulatory network inference can provide a broader context, but interpreting the complex connections within those networks can be challenging.

BLOBFISH simplifies the problem of subnetwork identification and interpretation by using gene regulatory network models to find a minimal set of genes and transcription factors linking the elements of a candidate gene set. As such, BLOBFISH complements more general network methods such as degree-based analyses, differential targeting analysis, and statistical comparisons of network edge weights; it fills an important methodological gap by extracting meaningful subnetworks in a way that balances the generalizability of genome-wide networks with the interpretability of a focused set of regulatory processes.

## Acknowledgements

This work was supported by the National Institutes of Health [R01HG011393, R35CA220523]. We thank Enakshi Saha, Kate Hoff-Shutta, and Kimberly Glass for support in the formulation of the null model for LIONESS networks and Viola Fanfani for support in data acquisition.

