## Supplementary Methods for "BLOBFISH: Bipartite Limited Subnetworks from Multiple Observations using Breadth-First Search with Constrained Hops"

### Null Model

We define the null network model as a model in which, by design, no biological signal is expected to be present. In the case of PANDA-like networks such as those used in our analysis, we constructed the null by retaining the known transcription factor binding motifs (motif) and protein-protein interactions (PPI) between transcription factors (which are constant across all samples) and generating ten random gene expression networks, where the expression is drawn from a normal distribution. In these random gene expression networks, the expression levels of any two genes are generated independently; therefore, they are expected, on average, to contain only uncorrelated or spuriously correlated genes. Each random gene expression network is then input into PANDA along with the motif and PPI. After running PANDA, the edges comprising the motif and PPI were removed from the final networks, and the null distribution of edge weights was represented as the distribution of edge weights remaining in the final networks.

### Pseudocode

The BLOBFISH algorithm is described as pseudocode below, where *n* is the number of observations, *S* is an aggregate weighted adjacency list with $\left| E \right|$ rows and 2 + *n* columns, where columns 1 and 2 represent source and target nodes and columns 3 through 2 + *n* represent weights, *G* is the seed set, *α* is the significance cutoff for statistical testing, ∅ is a set of edge weights obtained from the null model, and *h* is the maximum number of hops allowed to connect a pair of genes in *G*.

```
function BLOBFISH(S,G,α, Ø,h):
   SphereOfInfluence := # List of lists of edge sets, with
                        #dimensions |G| and h/2
   // Find all significant edges.
   SigEdges := []
   For edge in 1..|E|:
      If FDR(pval(Wilcox.test(S[e, 3:n+2], Ø)) < α:
         SigEdges += e

   // Obtain the sphere of influence for each starting gene.
   For g in G:
	Start := g
Explored := []
For i in 1..(h/2): # Ranging from 1 to h/2
   SphereOfInfluence[g][i]:= BFS(S,Start)
```

Explored := Explored ∪ Start

```
   Start := SphereOfInfluence – Explored

   // Find all paths connecting genes’ sphere of influence.
   Subnet := []
   For g in G:
	For γ in G – g:
	   For i in 1..(h/2):
```

Subnet += SigSubnet[g][i] ∩ SigSubnet[γ][i]

return Subnet

### Theoretical Analysis of Runtime from Pseudocode

**Lines 1-4:** The initialization of SigEdges and SphereOfInfluence are constant-time operations $O\left( 1 \right)$.

**Lines 5-7:** All statements in the loop For edge in 1..|E|:run in $O\left( n\log n \right)$, the time to sort the null and test distributions when computing a Wilcoxon rank-sum test. Therefore, the total running time of the loop is $O\left( n\log\left( n \right)|E| \right)$

**Lines 10-16:** All statements in the loop g in G: run in the amount of time required to run the loop on lines 13-16 (For i in 1..(h/2):) because the initialization of Start and Explored are constant time operations. Union and difference operations are at most linear in the number of total elements, which in this case is at most the number of vertices in *S* $\left| V \right|$ in addition to the number of vertices in *S* not belonging to the current node type (i.e., source or target), $\left| V_{2} \right|+\left| V \right|<2\left| V \right|$. Equivalence will occur iff SphereOfInfluence for a node includes all nodes not belonging to the current node type ($V_{2}$)and Explored contains all nodes $\left| V \right|$. Therefore, the runtime of lines 14-16 is $O\left( \left| V \right| \right)$, making the runtime of lines 13-16 $O\left( h\left| V \right| \right)$ and the total runtime of lines 10-16 $O\left( h\left| G \right|\left| V \right| \right)$*.*

**Lines 19-24:** Line 19 is a constant time operation. All statements in the loop g in G: run in the amount of time required to run the loop on lines 21-23 (For γ in G – g:). All statements in this loop run in the amount of time required to run the loop on lines 22-23 (For i in 1..(h/2):). The intersection operation runs in at most linear time, which is at most twice the number of vertices in *S*, $2\left| V \right|$ (with equivalence occurring when each gene’s sphere of influence at hop i includes all vertices not belonging to the current node type). Therefore, the runtime of lines 22-23 is $O\left( h\left| V \right| \right)$, the runtime of lines 21-23 is $O\left( h\left| G \right|\left| V \right| \right)$, and the runtime of lines 19-24 is $O\left( h\left| G \right|^{2}\left| V \right| \right)$*.*

The running time is therefore $O\left( max\left( h\left| G \right|^{2}\left| V \right|,h\left| G \right|\left| V \right|,n\log\left( n \right)|E| \right) \right)$, which is $O\left( max\left( h\left| G \right|^{2}\left| V \right|,n\log\left( n \right)|E| \right) \right)$, which is $O\left( max\left( h\left| G \right|^{2}\left| V \right|,n\log\left( n \right){|V|}^{2} \right) \right)$, which is $O\left( max\left( h\left| V \right|^{3},n\log\left( n \right){|V|}^{2} \right) \right)$, because $\left| G \right|$ is bounded above by $|V|$. In general, $n\log\left( n \right)\ll|V|$ for any whole-omics network. Therefore, the running time is $O\left( h\left| V \right|^{3} \right)$, which is polynomial in $\left| V \right|$.

### Theorem 1

Any path $P_{b,c}$ between nodes *b* and *c* in *G* contains a node *a* such that $D_{a,b}=D_{a,c}$, where $D_{i,j}$ represents the shortest unweighted graph distance between two nodes *i* and *j*.

**Proof:**

Because *b* and *c* are of the same node type in a bipartite network, $P_{b,c}$ must contain an odd number of intermediate nodes and an even number of connecting edges, i.e., $mod\left( D_{b,c},2 \right)=0$.

Further, we have that $D_{b,c}=D_{c,b}$.

Then, traversal along the first $\frac{D_{b,c}}{2}$ edges in $P_{b,c}$ starting at *b* results in a path $P_{b,x}$where $x\notin\left\{ a,b \right\}$.

Further, backwards traversal along the last $\frac{D_{b,c}}{2}$ edges in $P_{b,c}$ starting at *c* results in a path $P_{c,y}$where $y\notin\left\{ a,b \right\}$.

Therefore, $D_{b,x}=D_{c,y}=\frac{D_{b,c}}{2}$. If $x=y$, then *x* satisfies the condition for node *a*.

We can then assume for purposes of contradiction that $x\neq y$.

Then either *x* is not connected to *y* or $D_{x,y}>0$.

Because $P_{b,c}$ contains both *x* and *y*, not all edges in $P_{b,c}$ are connected, contradicting its definition as a path.

If $D_{x,y}>0$, then $D_{b,c}=D_{b,x}+D_{x,y}+D_{c,y}>D_{b,x}+D_{c,y}$.

Substitution yields $D_{b,c}>\frac{D_{b,c}}{2}+\frac{D_{b,c}}{2}$, contradicting the axiom of equality, i.e. that $D_{b,c}=D_{b,c}$.

Therefore, by contradiction, $x=y$.

Therefore, $x=a$, proving that $P_{b,c}$ contains a node *a* such that $D_{a,b}=D_{a,c}$,
