## Supplementary Figures for "BLOBFISH: Bipartite Limited Subnetworks from Multiple Observations using Breadth-First Search with Constrained Hops"

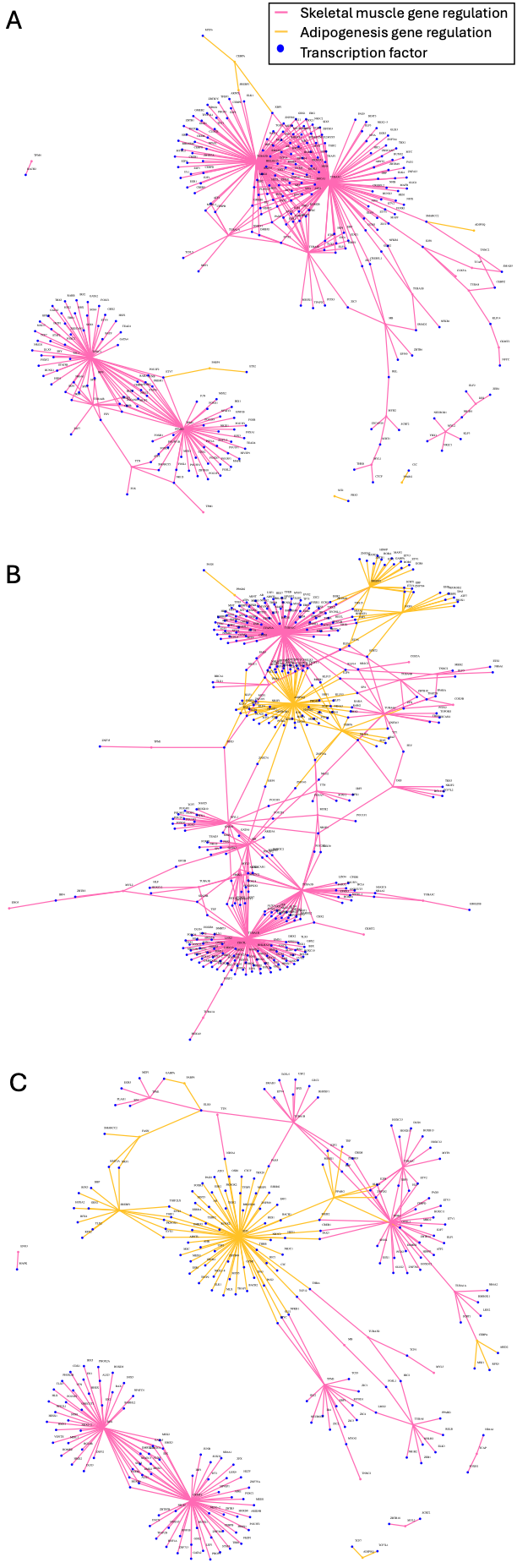

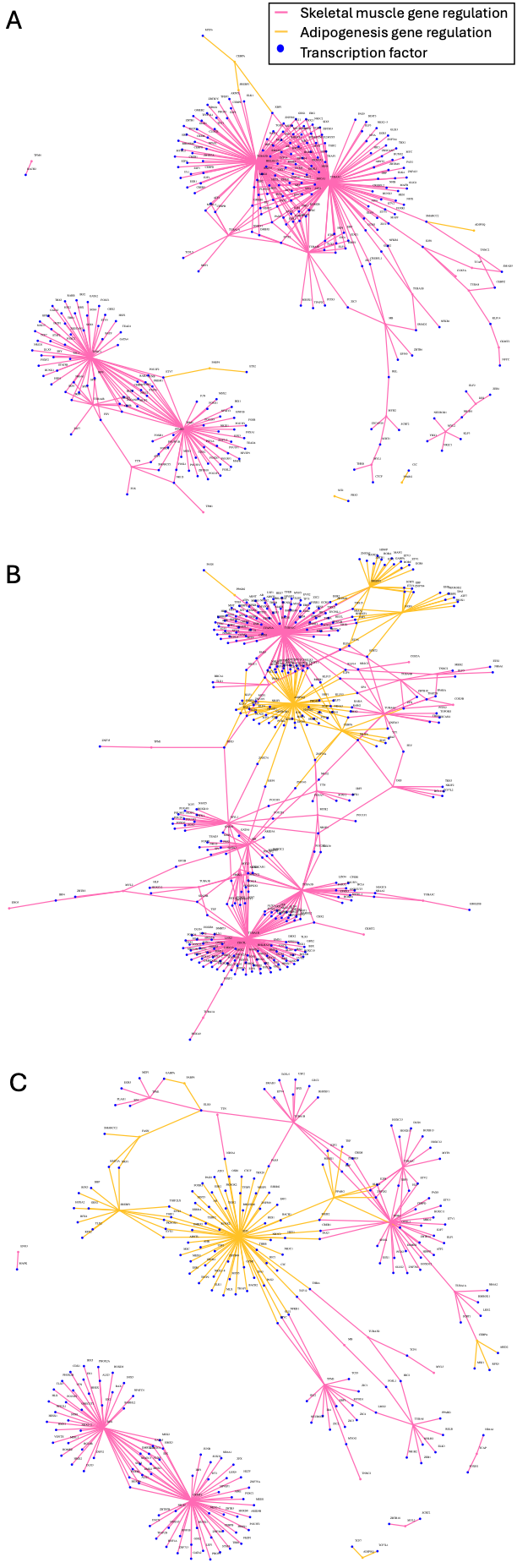

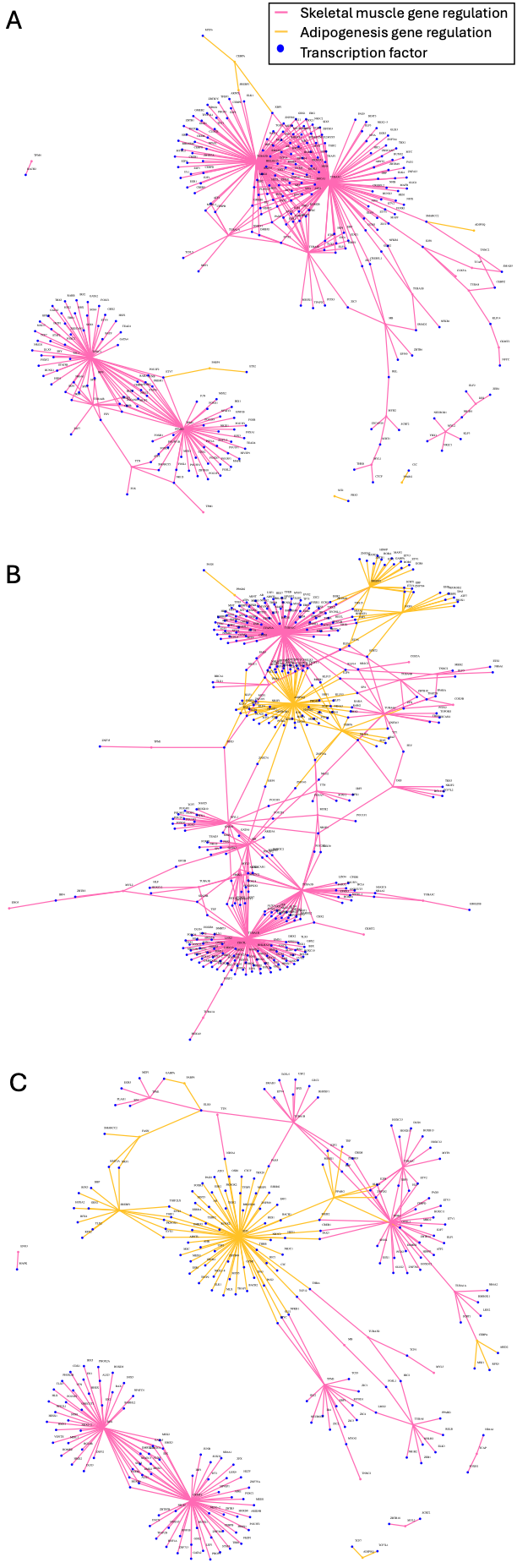

Supplementary Figure 1. Tissue-specific gene regulatory subnetworks for GTEx skeletal muscle samples for males 20-29 in (A) lung tissue, (B) skin, and (C) aorta.
